## Supplementary Material for "Predictions enable top-down pattern separation in the macaque face-processing hierarchy"

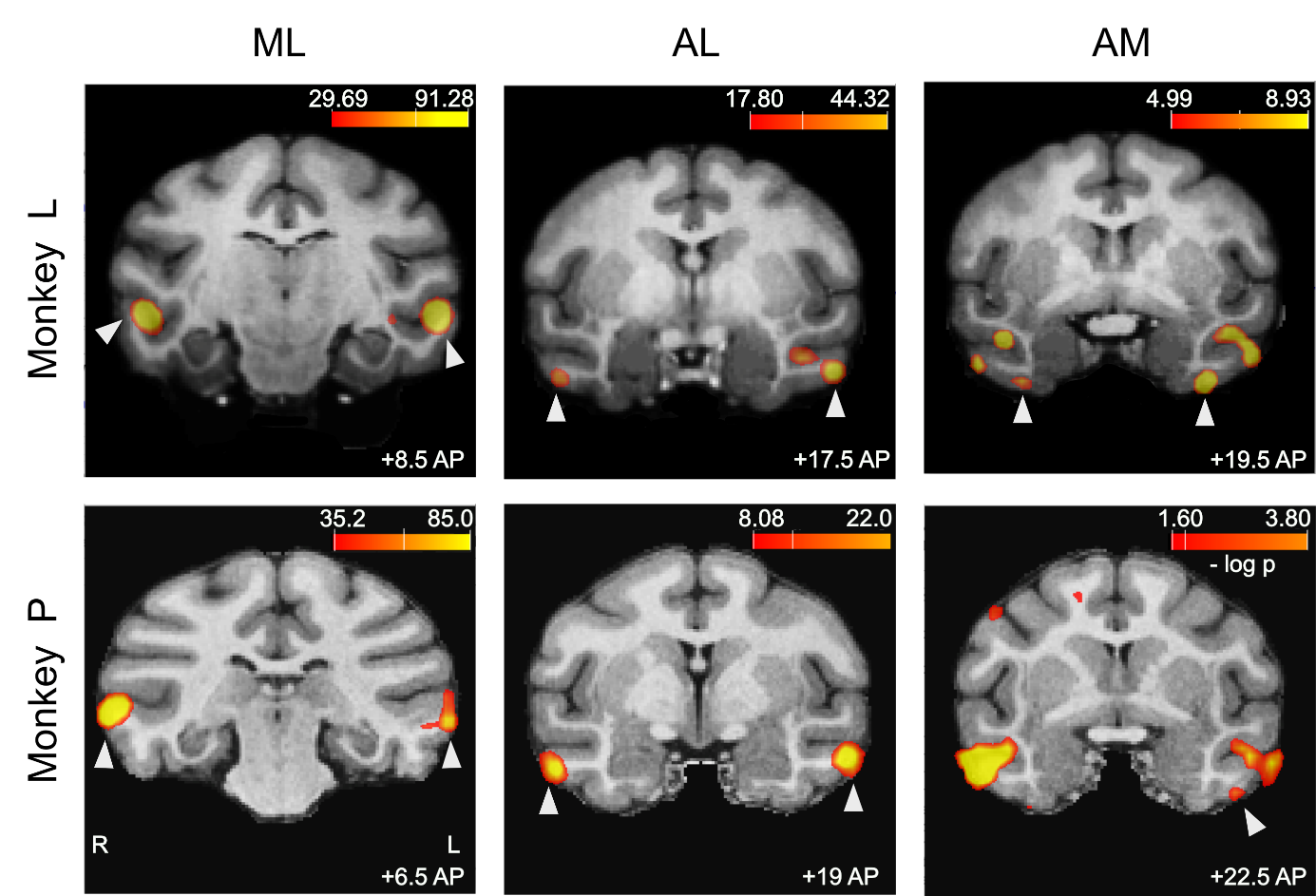

### Supplemental Information

**Figure S1:** **Face-areas localized in both monkeys using functional MRI**

Significance maps from the contrast: [faces vs. all other categories] are overlaid on coronal slices of anatomical MRIs. All localized face-areas are indicated with a white arrow (AM in the right hemisphere of Monkey P is not shown on this slice). Color bars indicate the negative log p-values of the significance map. Positive values of AP (in mm) indicate anterior from the ear-bars.

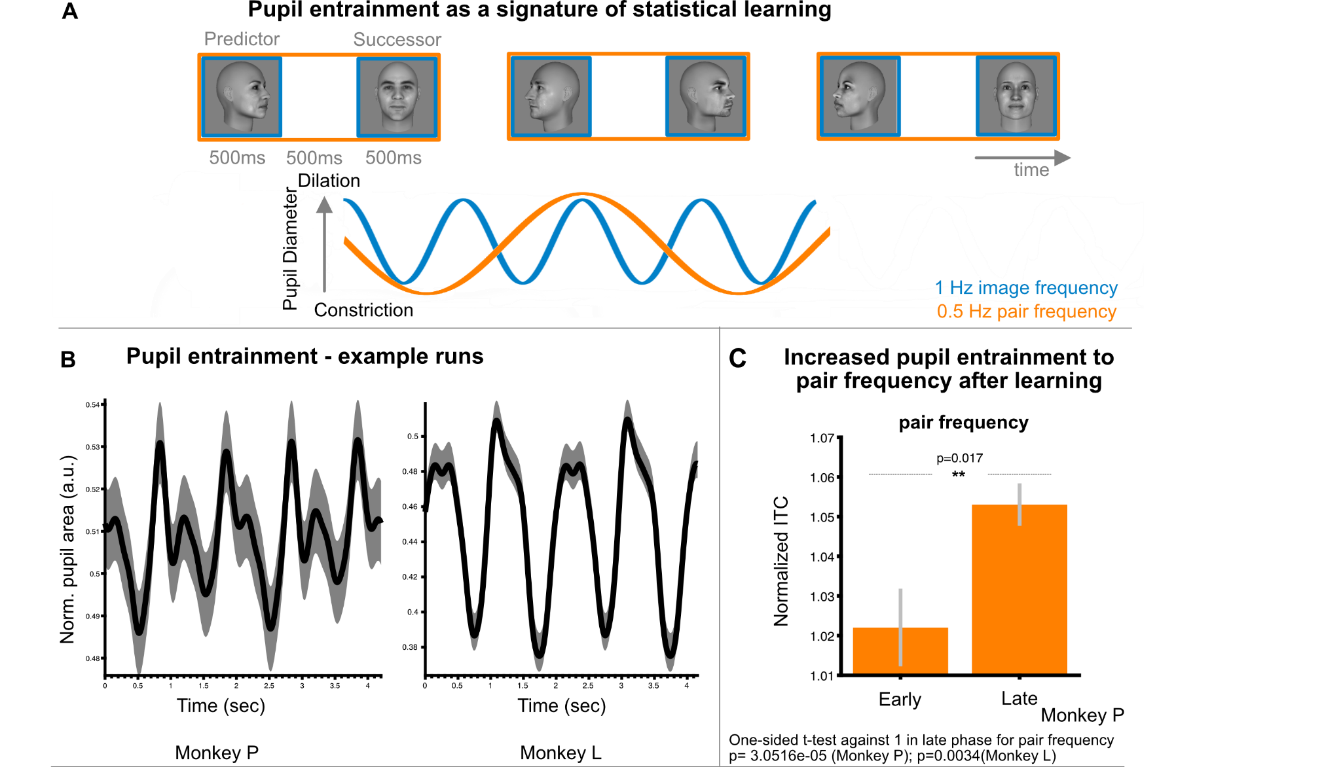

**Figure S2: Pupil entrainment as a signature of statistical learning of face-pairs.
A.** We presented a continuous sequence of face images at a rate of 1 Hz (image duration 500 ms, inter-stimulus interval 500 ms). The faces were systematically arranged such that one specific image predicted one other image, forming pairs. Hence, the frequency at which the pairs were presented was 0.5 Hz. The pupil is known to entrain to the image frequency (at 1 Hz) and to the pair frequency (at 0.5 Hz) once the pair structure has been learned^1^. **B.** Example runs showing pupil dilation dynamics to the face-pairs for both monkeys. The plots show averaged normalized pupil area (a.u.) aligned to the start of each pair. Shading indicates SEM. **C.** We found significant phase locking, quantified by inter-trial phase coherence (ITC), at the pair frequency (0.5 Hz) in both monkeys (Monkey P: p=3.0561e-05; Monkey L: p=0.0034, one-sided t-test against 1). ITC at the pair frequency increased after learning (p=0.017, unpaired one-sided t-test; data only available for Monkey P). Error bars depict SEM.

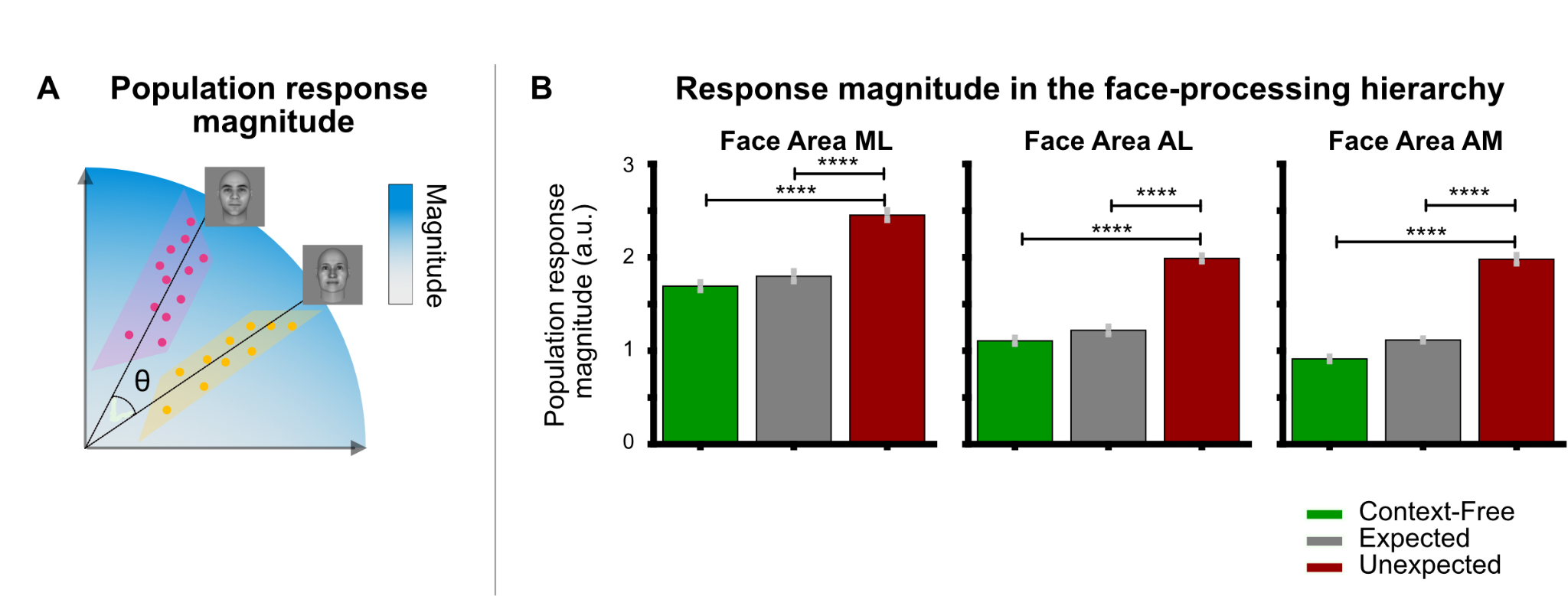

**Figure S3: Population response magnitude per condition and face-area. A.** Population response magnitudes were quantified to distinguish differences in the magnitude of the IT population response from angular differences in stimulus identity. Population response magnitude and population response vector direction are thought to contain differential information^2,3^. Response magnitude was quantified as the Euclidean (L2) norm across all voxels in a given ROI for each stimulus separately and then averaged. There was no significant correlation between the population magnitude response and pattern-separability (vector direction) across areas and conditions (Spearman’s rho=0.0167, p=0.9816). **B.** Prediction errors (PEs) are often described in terms of the magnitude of neural activity. We found higher response magnitudes when the same faces appeared as a violation of an expectation (red, unexpected condition) than when they were expected (gray) in all face-areas (paired t-tests, all p<3.5e-04, corrected for multiple comparisons). Response magnitudes were also larger in the unexpected than in the context-free condition (green; unpaired t-tests, all p<1.5e-07, corrected for multiple comparisons). Error bars depict SEM.

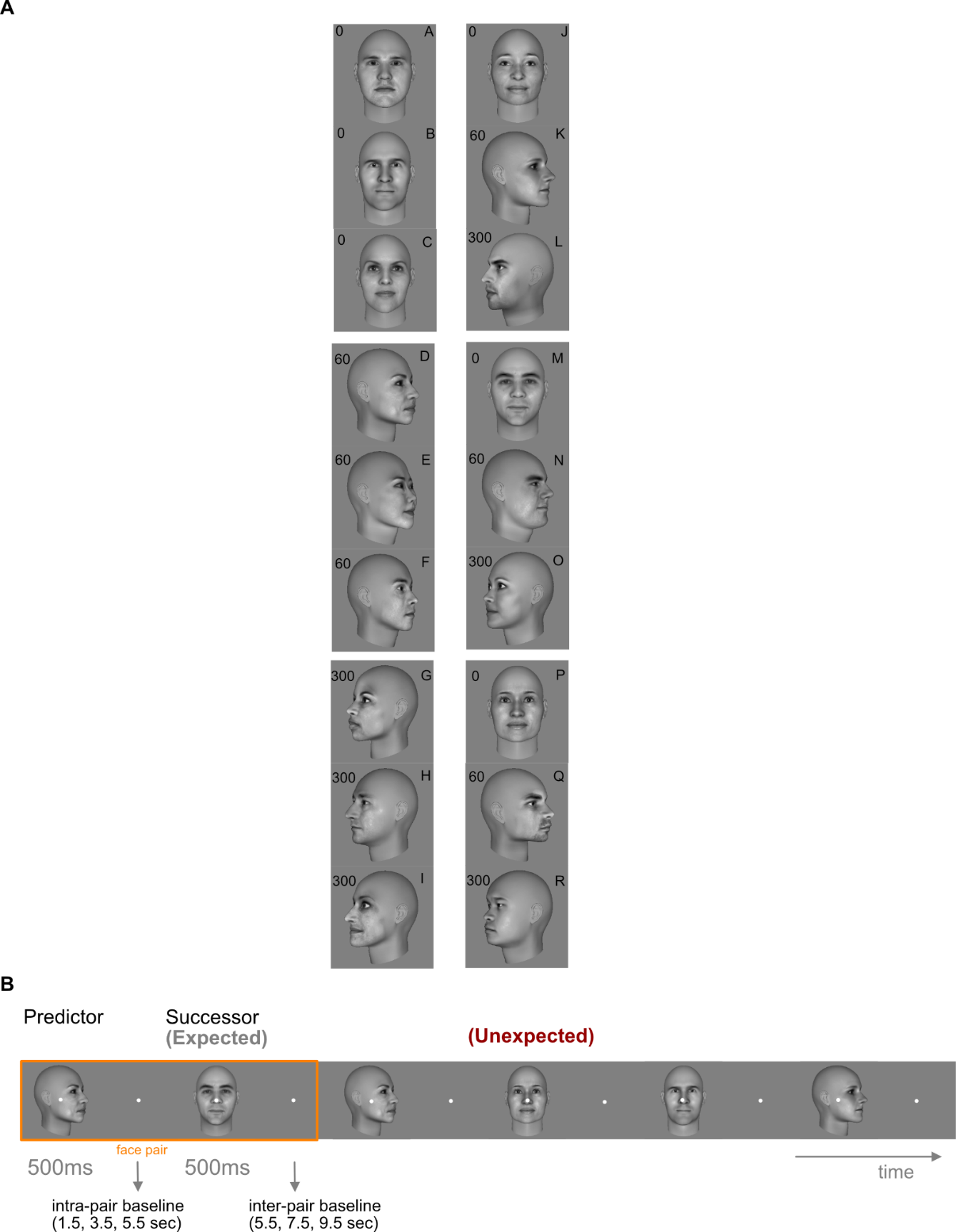

**Figure S4: Stimuli and task-schematic. A.** Trained Pairs in the statistical learning paradigm: In the training phase of the statistical learning paradigm, the monkeys were exposed to face-pairs to establish associations between the predictor and the successor images. Pairs were arranged such that a predictor face (one identity-view combination) would uniquely predict its specific successor face (one other identity-view combination). Training was conducted with sequentially presented face-pairs for at least 3 weeks for both monkeys. The sequence of pairs was designed such that the transitional probability within a pair was 100% and the transitional probability between pairs was kept at a minimum. **B.** Task-design for the test phase: In the test phase, monkeys were presented with a sequence of faces in a rapid event-related design during fMRI. Each face image was shown for 500 ms followed by a baseline period and each predictor face was followed by a successor face. The baseline period was temporally jittered to be able to extract single-trial responses from the BOLD signal. Baseline periods were pseudo-randomly selected (intra-pair baseline: 1.5, 3.5, or 5.5 s; inter-pair baseline: 5.5, 7.5, or 9.5 s). Trained pairs, i.e., ‘expected’ successors, were presented in 60% of all trials. Prediction violations were introduced by changing the successor image.

|  | | **Model Fits (Z-Spearman correlation)^a^** | | | | **p-values (bootstrap test; one-sided)^b^** | | | |
| --- | --- | --- | --- | --- | --- | --- | --- | --- | --- |
|  |  | Shape | Mirror-Symmetry | Appearance | View-Independent Appearance | Shape | Mirror-Symmetry | Appearance | View-Independent Appearance |
| **Context-Free** | ML | 0.3219 | -0.0263 | 0.1950 | -0.1990 | 0.0253 | 0.5610 | 0.1410 | 0.8598 |
|  | AL | -0.3425 | 0.6244 | -0.3469 | -0.2425 | 0.9732 | 9.9999e-5 | 0.9738 | 0.9178 |
|  | AM | 0.1615 | -0.1990 | 0.4740 | 0.5277 | 0.1717 | 0.8940 | 0.0030 | 0.0015 |
| **Expected** | ML | -0.1057 | 0.3437 | 0.0477 | 0.0872 | 0.7322 | 0.0213 | 0.4008 | 0.3279 |
|  | AL | -0.0039 | -0.0010 | 0.2606 | 0.8136 | 0.5106 | 0.5049 | 0.0710 | <0.0001 |
|  | AM | -0.2758 | 0.0371 | -0.1181 | -0.1216 | 0.9315 | 0.4304 | 0.7636 | 0.7574 |
| **Prediction Error** | ML | -0.0776 | 0.1795 | -0.0891 | 0.0934 | 0.6596 | 0.1574 | 0.6902 | 0.3016 |
|  | AL | 0.0686 | -0.2619 | 0.2425 | 0.3538 | 0.3450 | 0.9399 | 0.0953 | 0.0283 |
|  | AM | -0.1686 | 0.3207 | -0.0508 | -0.0986 | 0.8492 | 0.0257 | 0.6213 | 0.7129 |

**Table S1: Model-fits in representational similarity analysis** (Fisher-transformed Spearman rank correlation coefficients).

^a^ 2^nd^ level similarity between the models: shape, mirror-symmetry and appearance to the experimental 1st level RDMs was calculated as Spearman rank correlation while partialling out low-level similarities^4^ using a Gabor wavelet pyramid as a model of early visual cortex. 2^nd^ level similarity for the model view-independent appearance was calculated as Spearman rank correlation.

^b^ uncorrected p-values.

### Supplementary References

1.	Schwiedrzik, C.M., and Sudmann, S.S. (2020). Pupil diameter tracks statistical structure in the environment to increase visual sensitivity. J. Neurosci., JN-RM-0216-20. 10.1523/JNEUROSCI.0216-20.2020.

2.	Rust, N.C., and Cohen, M.R. (2022). Priority coding in the visual system. Nat. Rev. Neurosci. *23*, 376–388. 10.1038/s41583-022-00582-9.

3.	Jaegle, A., Mehrpour, V., Mohsenzadeh, Y., Meyer, T., Oliva, A., and Rust, N. (2019). Population response magnitude variation in inferotemporal cortex predicts image memorability. eLife *8*, e47596. 10.7554/eLife.47596.

4.	Kietzmann, T.C., Swisher, J.D., König, P., and Tong, F. (2012). Prevalence of Selectivity for Mirror-Symmetric Views of Faces in the Ventral and Dorsal Visual Pathways. J. Neurosci. *32*, 11763. 10.1523/JNEUROSCI.0126-12.2012.
